# DeepTracer: Predicting Backbone Atomic Structure from High Resolution Cryo-EM Density Maps of Protein Complexes

**DOI:** 10.1101/2020.02.12.946772

**Authors:** Jonas Pfab, Dong Si

**Affiliations:** Division of Computing and Software Systems, University of Washington, Bothell, USA

## Abstract

**Motivation:** Accurately determining the atomic structure of proteins represents a fundamental problem in the field of structural bioinformatics. A solution would be significant as protein structure information could be utilized in the medical field, e.g. in the development of vaccines for new viruses. This paper focuses on predicting the protein structure based on 3D images of the proteins captured through cryogenic electron microscopes (cryo-EM). A fully automated computationally efficient protein structure prediction method would be particularly beneficial in the field of cryo-EM as the technology allows researchers to photograph multiple large protein complexes in a single study, which means that a fast prediction method could allow for a high throughput of derived protein structures. We present a deep learning approach, DeepTracer, for predicting locations of the backbone atoms, secondary structure elements, and the amino acid types. In order to connect the predicted amino acids into chains, we applied a modified traveling salesman algorithm.

**Results:** We trained our deep learning model on experimental cryo-EM density maps and tested it on a set of 50 density maps. We found that our new approach predicted protein structures with an average RMSD value of 1.18 and a coverage of 87.5%. Furthermore, we detected secondary structure information for 87.2% of amino acids correctly. We also showed preliminarily that 25.2% of amino acid types could be predicted directly from the 3D cryo-EM density map, considering 20 different types in total. Finally, we noted that the prediction runtime of DeepTracer is significantly improved compared to other methods. It predicts a large protein complex structure of more than 30,000 amino acids in only 2 hours.

**Availability:** The repository of this project will be published.

**Supplementary information:** Supplementary data will be available at *Bioinformatics* online.

## 1 Introduction

Proteins represent an essential building block in all organisms. They perform crucial functions such as DNA replication, responding to stimuli, and many more [1]. Each protein consists of a sequence of the same 20 kinds of amino acids, while all amino acids are made up of a carbon, carbon-alpha (Cα), and nitrogen atom, as well as a side chain which determines the amino acid type and is connected to the Cα atom. The functionality of a protein is determined through its structure, defined by the three-dimensional (3D) arrangement of the amino acids that make up the protein [2].

As a consequence, knowledge about the structure of a protein helps to gain a deeper understanding about its behavior. This can be particularly useful, e.g., in the development of new vaccines and drugs for new viruses such as Novel Coronavirus (2019-nCoV) [3]–[5], as the surface proteins of a virus determine how it will interact with a person [6], [7]. Therefore, it is fundamentally important to be able to accurately predict the 3D structure of a protein.

There are two main approaches to predict protein structures: homology modeling and de novo structure prediction. They differ in such that homology modeling requires structural knowledge about a homologous protein for its prediction while de novo methods do not. This limits the application range of homology modeling, since homologues proteins must be obtained first before making predictions, which is why we aim at the more challenging de novo structure prediction. Traditionally, research on de novo protein structure prediction was focused on inferring the 3D structure through the protein’s amino acid sequence [8]. This approach is beneficial as the determination of the sequence is relatively straightforward. Methods that follow this approach either use geometric calculations to estimate the protein structure [9], [10], or they use deep learning techniques, such as Google’s AlphaFold [11]. While these methodologies have led to impressive results, their accuracy is naturally limited as the amino acid sequence does not completely contain the protein’s structural information, especially for very large protein complexes [12].

Throughout the last decade, technological advancements in cryogenic electron microscopy (cryo-EM) have provided a new approach to solve the structure determination problem. Cryo-EM allows researchers to capture 3D images of large protein complexes by cooling them to cryogenic temperatures without the need of a costly crystallization of the protein [13]. In recent years the resolution of these images has increased to a near-atomic level which led to an exponential gain in popularity of the technology [14]. A 3D image captured through cryo-EM is an electron density map (see Fig 1). It represents the volume of the protein complex through a 3D grid of voxels. Each voxel stores the electron density value, meaning the probability that an electron is present, for its location. Latest developments in the field of cryo-EM have allowed researchers to capture many high-resolution density maps from large protein complexes in the course of a single study [15]. This makes the ability to predict protein structures from cryo-EM maps fully automatically even more significant as it allows for a very high throughput of protein structures derived from cryo-EM maps.

**Fig 1.**
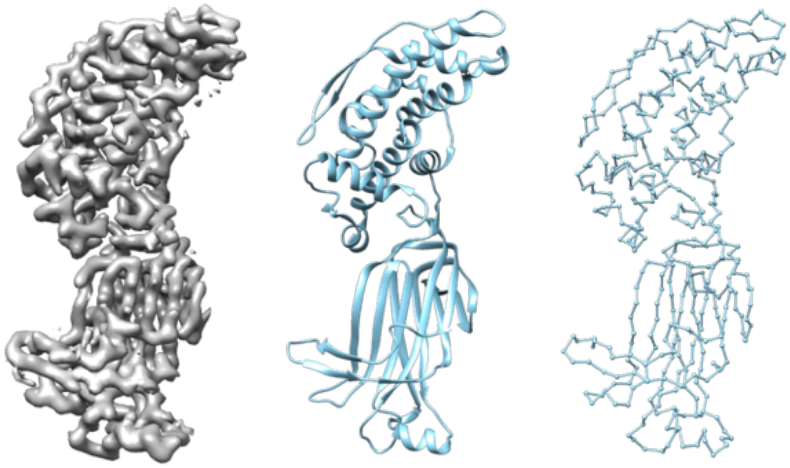
EMD-6272 (rotavirus VP6) Density Map (left) and its corresponding solved structure PDB-3j9s depicted in ribbon (center) and backbone view (right).

The structure of a protein can be estimated from its density map, intuitively by placing the amino acids in such way that the resulting structure fits well into the density map. While this sounds like a simple task it becomes very challenging due to lower local resolutions as well as noise created through errors in the cryo-EM. Currently, there are several existing methods that perform protein structure predictions based on cryo-EM data such as Phenix and Rosetta [16], [17]. They both use conventional algorithms to place and connect the amino acids of the protein. However, they often struggle to identify amino acids as such and, therefore, tend to have a low coverage on large protein complex [18]. Furthermore, their long runtime is a problem. The prediction of small density maps can take up to several days rendering the method practically useless for the prediction of large proteins. The C-CNN method, another protein structure prediction application which utilizes deep learning techniques, achieves better results in terms of coverage [18]. However, it requires some manual input and the runtime could be further improved. Additionally, it predicts the location of Cα atoms without information about their amino acid type. Therefore, we present the DeepTracer, a new fully automated method for the prediction of protein structures from cryo-EM density maps, with the goal of improving the runtime by applying a 3D U-Net in combination with a modified travelling salesman algorithm and having the ability of recognizing the amino acid type from 3D cryo-EM density maps.

The functionality of our new DeepTracer is explained in the Methods section. In the Results section, the DeepTracer is applied on a test set of 30 density maps in order to assess and compare its accuracy with other methods. The implications of the project and possible future work is examined in Discussion. Finally, we recapture our findings in the Conclusion section.

## 2 Methods

The full prediction pipeline of the DeepTracer includes many different steps. In this section we will look at the major ones and explain how they create a protein structure based of a density map. We begin by looking at the 3D U-Net which predicts confidence maps that show where e.g. atoms might be or where which secondary structure is likely to be present. Next, we will examine how the DeepTracer uses this information to place the actual amino acids, as well as how it connects them into chains to form the protein structure. Finally, we discuss how secondary structure information can be used to refine predicted α-helices.

### A. 3D U-Net Deep Learning Model

The deep learning model represents the central entity in the structure prediction. Its job is to predict not only the locations of amino acids but also secondary structure positions and amino acid types. To explain its realization, we start by looking at its architecture. Then, we move on to the data collection and preprocessing steps and finally describe the model’s training process.

The U-Net model gets its name from the U-shape of its architecture which is optimized for segmentation tasks on medical images [19]. The detailed architecture of the model can be seen in **Fig 2**. The cryo-EM density maps are fed to the 64^3^ input layer. The output layer has the same 64^3^ shape with N different channels. The number of channels depends hereby on what the U-Net is predicting (e.g. the secondary structure U-Net has three channels for loops, sheets, and helices).

**Fig 2.**
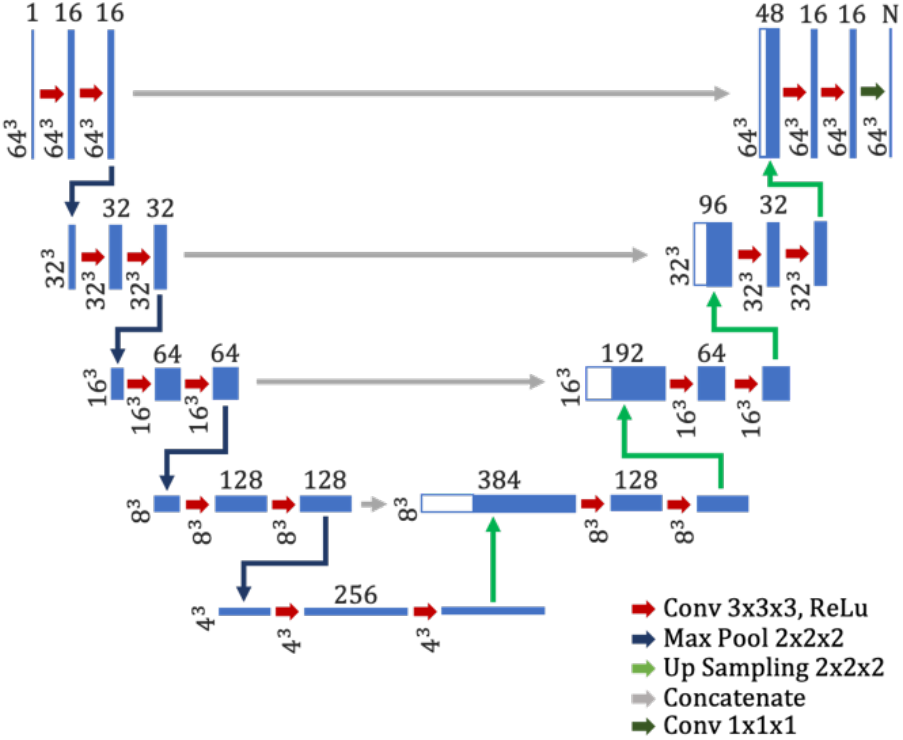
Architecture of the U-Net as used by the DeepTracer. The blue boxes show the output maps of the different layers where the dimensions of the maps are depicted on the left and the number of channels is depicted on top.

As mentioned above, the U-Net should predict multiple things, including the amino acid positions, secondary structure elements, as well as the amino acid types. Therefore, the model consists of three parallel U-Nets as shown in **Fig 3**. The first one is responsible for predicting the amino acid positions. As the Cα atom of each amino acid is central to its position we can reduce this task to only predicting the Cα atoms of the protein structure. Additionally, this U-Net also predicts the backbone location of the protein, which is defined as the position of all Cα, C, and N atoms of the structure. This prediction will later on be useful in order to connect the amino acids into chains. Thus, the output of the Cα atoms U-Net has two channels. The second U-Net is responsible for predicting the secondary structure elements and, therefore, its output has three channels, one for each structural element (α-helix, loop, and sheet). Finally, the third U-Net is tasked to predict the amino acid types of the protein structure. Since there are 20 different types of amino acids occurring in nature its output has the same number of channels. In **Fig 4** we can see an example output for each of the U-Nets with the density map from **Fig 1** as input.

**Fig 3.**
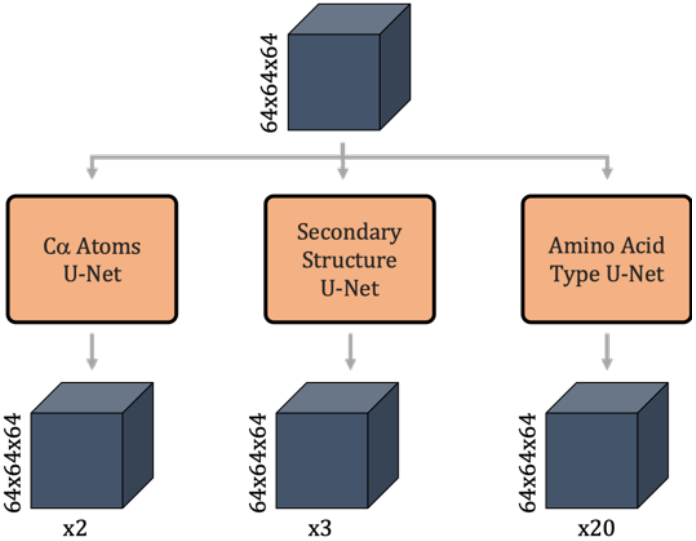
Architecture of the deep learning model, which includes three parallel U-Nets. The blue boxes show the input and output maps of the model, with their dimensions noted to the left and the number of channels marked below.

**Fig 4.**
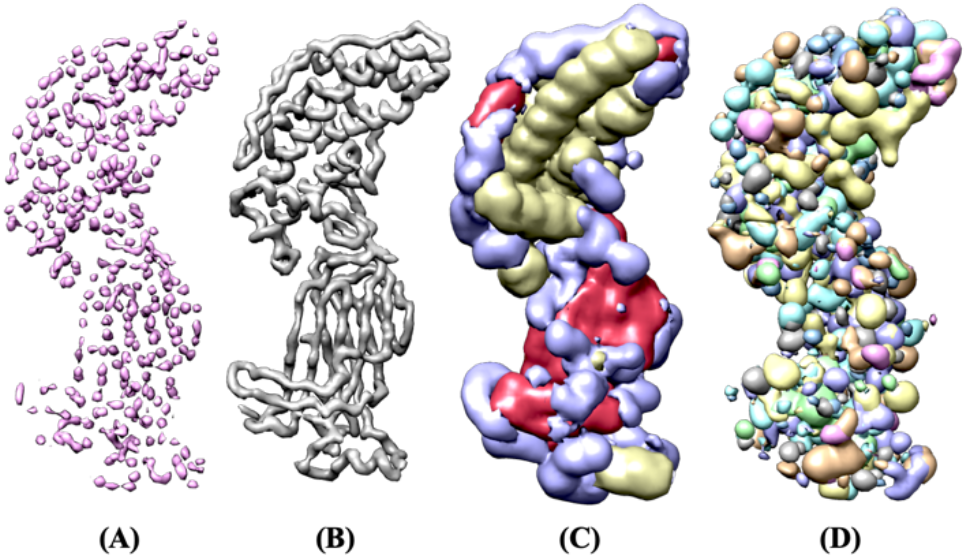
Example output confidence maps of the model for the EMD-6272 density map. (A) and (B) are both the output of the Cα atom U-Net and show the Cα atom locations (A) as well as the backbone of the protein (B). (C) shows the output of the secondary structure U-Net with a-helices in yellow, loops in purple, and sheets in red. In (D) we can see the output of the amino acid type U-Net where each color represents a different type of amino acid.

Now that we have looked at the architecture of the model, we can continue with the data collection. In order for the deep learning model to learn common noises and errors present in cryo-EM density maps we decided to train the U-Net using experimental data rather than density maps simulated from solved structures such as [18]. We downloaded the density maps from the EMDataResource website [20] in combination with their solved protein structures which function as the ground truth for the training process. We obtained the solved structures from the RCSB Protein Data Bank [21]. Since this paper is focused on high resolution maps, we only used density maps with a resolution of 4Å or better. In total we used 411 different density maps to build the training and validation sets.

The preprocessing steps can be split into preparing the cryo-EM density maps and creating the different labels based on the solved structures. For the experimental density maps we have to make sure that the voxel size equals one, meaning that every voxel measures 1Å. Furthermore, we need to adjust the dimensions of the map to the 64^3^ input dimensions of the model as seen in Fig 2. This is done by splitting the maps into several 64^3^ big cubes. In order to avoid errors at the borders of the cubes we overlay them partly such that we only use the 50^3^ center voxels of each cube to reconstruct the output maps to the original dimensions of the density map. By splitting the 411 density maps that we collected in this manner, we retrieve 16,070 training examples. To create the labels that the model can be trained on, we use the solved structure to create masks by filtering out the corresponding atoms and setting all voxels within a certain distance to those atoms to one. For the Cα mask we filter only the Cα atoms of the protein structure and set all directly neighboring voxels around them to one. The backbone mask is created similarly except that we use all Cα, C, and N atoms. To generate the secondary structure masks, we filter out all atoms for each structural element and set all voxels within a distance of two around each filtered atom to one. Finally, to create the amino acid type masks we filter out all atoms for each amino acid type and set all neighboring voxels to one for each atom. In total this gives us 25 different masks for each training example (Cα mask, backbone mask, three secondary structure masks, and 20 amino acid type masks).

### B. Placing Cα atoms

As mentioned above, the determining factor for the position of an amino acid is the position of its Cα atom. In order to place the Cα atoms in the 3D space we use the Cα atoms confidence map as well as the backbone confidence map. The backbone map is used to identify different chains and the Cα atoms map to find the locations of the Cα atom.

Before we place any Cα atoms, we try to identify separate chains in the protein structure. This will help us later when connecting the atoms and improves the runtime of the connection process. In order to discern the chains, we identify connected areas of voxels that have a confidence value of 0.5 or higher. An example of the chains identified from a backbone prediction can be seen in **Fig 5**. We can now mask all other confidence maps with each chain area that we identified in order to continue the prediction independently for all chains.

**Fig 5.**
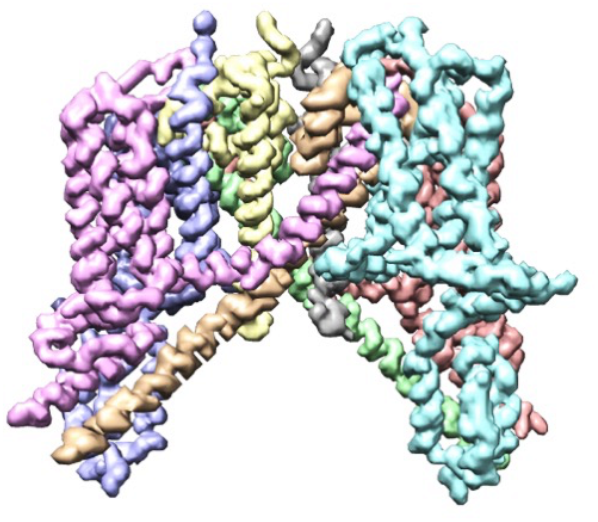
Backbone prediction of EMD-0478 density map (human TPC2 channel) (human TPC2 channel) with identified chains marked in different colors.

Next, we place the actual Cα atoms in two simple steps, using the masked Cα confidence maps. First, we find the initial atom locations by calculating local minima in the confidence map within a neighborhood of one voxel and a minimum value of 0.6. The value of 0.6 was chosen after some testing as it produced the most accurate results. This gives us a list of voxel indices. However, as atoms can have floating point coordinates, we try to improve this initial location by calculating the center of mass for 3^3^ cubes around each local minimum. The resulting values are used as the coordinates for the Cα atoms of the protein structure.

### C. Tracing Backbone

In this prediction step, we use the predicted Cα atoms and connect them into chains using a travelling salesman algorithm (TSP) [22]. However, instead of using the Euclidean distances between atoms to measure the total length of the path, we use a custom confidence score that expresses how confident we are that two atoms share a connection. The task of the TSP algorithm is then to connect the atoms into chains such that they maximize the sum of all confidence scores.

The calculation of the confidence score between two atoms takes two factors into consideration: The Euclidean distance between them, as well as the average confidence of voxels that lay in between the atoms in the backbone prediction. Let *p*(*x*, *μ*, *σ*) be the normal probability density function at *x* with mean *μ* and standard deviation *σ*, *d*_*a,b*_ the Euclidean distance between atoms a and b, and *b*_*a,b*_ the average backbone confidence between atoms a and b. We define the confidence score as shown in equation 1. The constant 3.8 was chosen as the mean for the distances between two atoms as this corresponds to the average distance found in nature. The standard deviation parameters of 5 and 0.3 where selected based on test runs. They were intentionally chosen higher than in the distribution found in solved structures, in order to cut some slack for prediction inaccuracies.

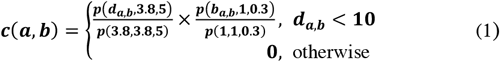

In order to apply the TSP algorithm, we also need to specify a start/end point. However, we do not know yet at which atom the chain will start and end. Therefore, we add a new atom that is connected to every other atom with a confidence of 1. This atom is then specified as the start/end and later on removed from the actual chain. An example of the application of the TSP on a list of Cα atoms can be seen in **Fig 6**.

**Fig 6.**
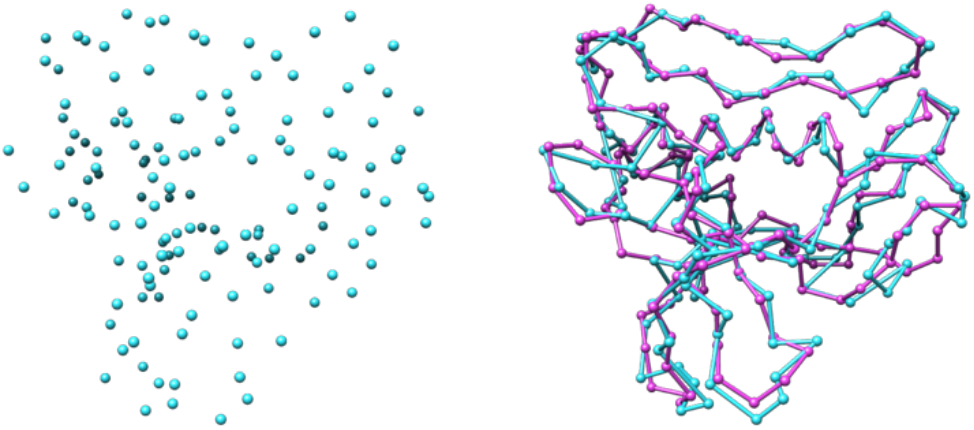
Predicted Cα atoms for EMD-4054 density map (major capsid protein) (major capsid protein) in blue before (left) and after (right) the backbone tracing step compared to the true structure in pink.

### D. Helix Refinement

The last prediction step is responsible for refining modelled α-helices. Here for, we can exploit the fact that the shape of an α-helix has a general definition which is valid across proteins [23]. Since the U-Net predicts the confidence of secondary structure elements, as shown in **Fig 4**, we know which amino acids belong to an α-helix based on the confidence of their region in space. We combine this knowledge of α-helix locations and their shape attributes in order to adjust the appropriate Cα atoms to better fit the shape of a natural α-helix structure.

For an α-helix which centers around the z-axis, we can use Equation 2 to model its shape where the variables *s* and *r* represent the initial shift and rotation of the helix. The values *2.11* and *1.149* are constants that define the radius and pitch of the helix to best match those of an α-helix.

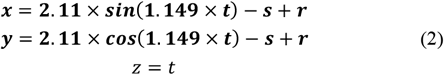

Equation 2 however, cannot be used to describe an α-helix which does not center around the z-axis or whose shape is not a straight cylinder. Since this is the case for most α-helices, it is necessary to adjust the equation in such way that it will address these issues. With the aim of doing so, we first locate the screw axis, the center line around which the helix winds itself, for each α-helix. This is achieved by calculating the centroid of consecutive intervals of the α-helix and then connecting them to approximate the true curve. An example of an α-helix and its calculated screw axis can be seen in Fig 7.

**Fig 7.**
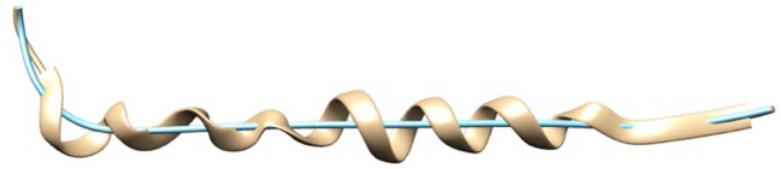
α-Helix extracted from the backbone prediction of the EMD-8515 density map in tan color and its screw axis in teal color.

Now that we know the location and shape of the screw axis for the α-helix, we need to incorporate this information into Equation 2. This is achieved by interpreting *t* as the distance that we travelled on the screw axis and use the unit direction vector of the screw axis at a certain point *t* as the new z axis. Next, we can find the new y-axis by calculating the cross product of the x-axis and the new z-axis and then normalizing it. Finally, we can get the new x-axis by calculating the cross product of the new z and y-axis and normalizing it again. By concatenating the three new axes we can get a rotation matrix **RM** with which we can calculate the point of the α-helix for any value *t* as shown in Equation 3.

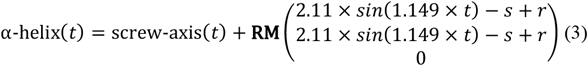

Now, we need to know the values *t* at which we have to insert Cα atoms. Since we know that an α-helix has a rise of 1.5Å per residue [23] we can increase *t* in steps of 1.5 and add a new Cα atom at α-helix(*t*).

In the final step we minimize the average distance from the Cα atoms of the refined α-helix to the Cα atoms of the original prediction. This is done by applying a minimization algorithm^1^ over the variables *s* and *r* to try different initial shifts and rotations. The final results of the α-helix refinement step are shown in **Fig 8**.

**Fig 8.**
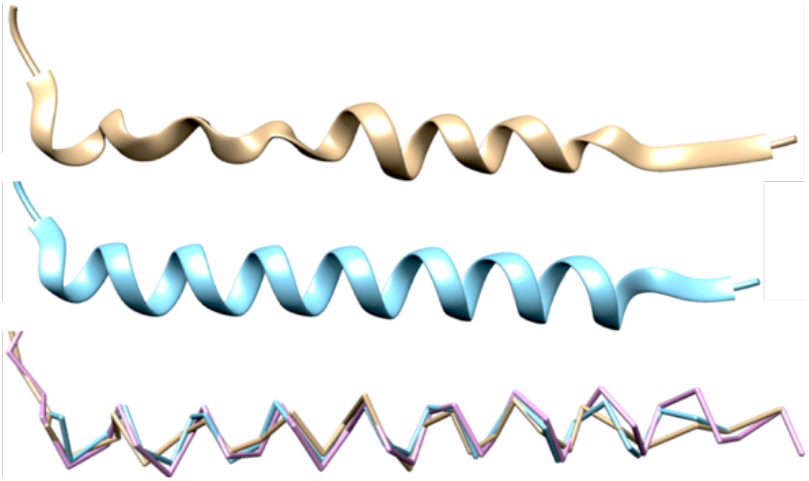
α-Helix extracted from the prediction of the EMD-8515 density map. (Top) Original prediction before the helix-refinement step. (Center) α-Helix after the refinement. (Bottom) Direct comparison of original prediction in tan color, refined prediction colored in teal, and the solved structure (PDB-5u70) in pink color.

## Results

In order to gain a better understanding about the performance of the DeepTracer we will apply it to a test set of 30 experimental density maps and evaluate the predictions using different metrics. Additionally, we will compare its performance against Phenix, as well as the C-CNN backbone prediction method.

We measure the accuracy of a predicted protein structure by comparing it to its corresponding solved structure using the following metrics. First, the root-mean-square deviation (RMSD) value describes the average distance between atoms of the predicted and solved protein structure. This gives us an idea how close the predicted atoms are to their correct counter parts. However, this does not give us any information about the percentage of atoms that we have correctly identified. For this purpose, we calculate the coverage value, which expresses the percentage of atoms in the solved structure that lay within 3Å of a predicted atom. Finally, we evaluate the false-positive percentage which is the number of atoms that were predicted further than 3Å from any atom in the solved structure, divided by the total number of predicted atoms. In addition to the metrics that we use to compare the other methods to the DeepTracer, we also calculate the secondary structure and amino acid type accuracy. We cannot evaluate these for the other methods as they utilize the U-Net confidence maps rather than the predicted protein structure. The secondary structure accuracy is defined as the percentage of atoms from the solved structure whose secondary structure type equals the one predicted by the U-Net for its location. The amino acid type accuracy is calculated in the same way. In Table 1, we can see the test results for all 30 density maps.

**Table 1.** Prediction results of the Phenix, C-CNN, and DeepTracer methods for 30 experimental density map.

| EMDB ID | PDB ID | Res. (Å) | Native<br>Ca<sup>2</sup> | FP %<sup>3</sup> |  | Coverage %<sup>4</sup> |  |  | RMSD |  |  | SSE %<sup>5</sup> | AA %<sup>6</sup> |
| --- | --- | --- | --- | --- | --- | --- | --- | --- | --- | --- | --- | --- | --- |
|  |  |  |  | C-CNN | Deep-Tracer | Phenix | C-CNN | Deep-Tracer | Phenix | C-CNN | Deep-Tracer | Deep-Tracer | Deep-Tracer |
| 2513 | 4ci0 | 3.4 | 893 | 3.5 | 3.6 | 62.5 | 96.6 | 91.5 | 1.33 | 1.06 | 1.13 | 84.99 | 35.05 |
| 3061 | 5a63 | 3.4 | 1223 | 3.0 | 5.3 | 73.9 | 95.6 | 91.0 | 1.15 | 1.10 | 1.14 | 90.02 | 29.44 |
| 3121 | 5aco | 4.4 | 2415 | 10.4 | 19.6 | 57.0 | 83.4 | 78.6 | 1.63 | 1.45 | 1.38 | 73.20 | 12.89 |
| 3222 | 5flu | 3.8 | 2119 | 1.1 | 4.7 | 62.0 | 86.6 | 82.0 | 1.76 | 1.60 | 1.56 | 70.60 | 8.82 |
| 3238 | 5fn3 | 4.1 | 1318 | 3.6 | 8.9 | 56.1 | 79.5 | 74.1 | 1.50 | 1.48 | 1.51 | 79.21 | 9.18 |
| 3601 | 5n8o | 3.9 | 1433 | 6.6 | 7.1 | 50.0 | 85.8 | 86.8 | 1.29 | 1.37 | 1.30 | 72.77 | 20.67 |
| 3785 | 5odv | 4.0 | 4968 | 10.0 | 12.4 | 40.1 | 91.5 | 89.8 | 1.73 | 1.46 | 1.40 | 82.77 | 21.36 |
| 4054 | 5lij | 4.2 | 152 | 1.3 | 1.4 | 84.2 | 90.8 | 88.2 | 1.44 | 1.44 | 1.28 | 71.05 | - |
| 4062 | 5ljv | 3.6 | 1950 | 3.8 | 2.9 | 48.3 | 95.8 | 91.9 | 1.21 | 1.10 | 1.06 | 81.85 | 33.95 |
| 4128 | 5lzp | 3.5 | 6182 | 0.8 | 2.9 | 77.1 | 93.2 | 88.9 | 1.13 | 1.16 | 1.17 | 87.45 | 27.71 |
| 5623 | 3j9i | 3.3 | 5978 | 0.8 | 2.1 | 72.4 | 95.9 | 92.0 | 0.87 | 1.05 | 0.92 | 87.77 | 36.95 |
| 5778 | 3j5p | 3.3 | 2368 | 1.2 | 5.1 | 55.6 | 74.7 | 68.1 | 1.15 | 1.29 | 1.49 | 77.58 | 17.27 |
| 5995 | 3j7h | 3.2 | 4088 | 1.5 | 1.9 | 72.9 | 96.9 | 93.4 | 1.12 | 1.10 | 0.99 | 82.85 | 29.55 |
| 6224 | 3j9c | 2.9 | 423 | 0.5 | 5.9 | 71.9 | 95.3 | 95.7 | 0.99 | 1.05 | 0.87 | 75.89 | 29.31 |
| 6239 | 3j9d | 3.3 | 746 | 1.9 | 2.4 | 74.8 | 98.1 | 94.8 | 0.99 | 1.01 | 1.02 | 78.28 | 34.18 |
| 6240 | 3j9e | 3.3 | 520 | 0.2 | 0.9 | 69.4 | 83.5 | 79.6 | 0.82 | 1.02 | 1.06 | 85.96 | 32.50 |
| 6272 | 3j9s | 2.6 | 397 | 0.0 | 2.5 | 88.9 | 99.0 | 97.0 | 0.69 | 0.85 | 0.87 | 88.16 | 31.49 |
| 6324 | 3ja7 | 3.6 | 5136 | 2.0 | 4.5 | 69.4 | 83.2 | 87.0 | 1.13 | 1.30 | 1.31 | 81.31 | 26.69 |
| 6346 | 3jaf | 3.8 | 1710 | 1.5 | 3.8 | 72.8 | 91.7 | 88.0 | 1.10 | 1.20 | 1.12 | 85.26 | 13.92 |
| 6408 | 3jb6 | 3.3 | 1731 | 1.3 | 2.6 | 79.3 | 96.6 | 92.5 | 0.83 | 0.97 | 1.11 | 88.85 | 36.45 |
| 6488 | 3jby | 3.7 | 1802 | 14.4 | 8.0 | 57.3 | 92.6 | 87.6 | 1.18 | 1.19 | 1.20 | 85.85 | 25.14 |
| 6551 | 3jcf | 3.8 | 1745 | 1.3 | 3.5 | 74.2 | 93.8 | 92.1 | 1.15 | 1.17 | 1.35 | 87.97 | 23.50 |
| 6630 | 3jcz | 3.3 | 2976 | 0.6 | 2.5 | 66.1 | 91.8 | 90.4 | 1.05 | 1.09 | 1.13 | 81.49 | 32.53 |
| 6631 | 3dj0 | 3.5 | 2976 | 1.9 | 2.3 | 59.5 | 89.4 | 88.8 | 1.07 | 1.16 | 1.19 | 80.44 | 25.44 |
| 6676 | 5wq8 | 3.3 | 7440 | 1.3 | 4.2 | 54.2 | 80.7 | 80.6 | 0.93 | 1.12 | 0.97 | 79.09 | 30.36 |
| 6703 | 5x58 | 3.2 | 3159 | 5.2 | 9.1 | 53.1 | 84.5 | 82.5 | 1.27 | 1.31 | 1.21 | 75.63 | 24.12 |
| 6744 | 5xno | 3.5 | 854 | 25.9 | 28.3 | 55.4 | 85.7 | 80.7 | 1.22 | 1.25 | 1.33 | 82.55 | 17.68 |
| 6770 | 5xsy | 4.0 | 1312 | 7.4 | 8.5 | 68.5 | 88.7 | 88.0 | 1.21 | 1.32 | 1.41 | 86.81 | 20.96 |
| 7063 | 6b7n | 3.3 | 2898 | 4.1 | 7.2 | 71.3 | 96.4 | 89.6 | 1.21 | 1.14 | 1.08 | 81.19 | 18.12 |
| 8015 | 5gaq | 3.1 | 2592 | 0.5 | 1.1 | 79.2 | 99.0 | 93.2 | 1.10 | 0.91 | 0.77 | 84.84 | 25.54 |
| Avg. |  |  |  | 3.9 | 5.8 | 67.1 | 90.5 | 87.5 | 1.10 | 1.19 | 1.18 | 81.72 | 25.20 |

<sup>2</sup> Number of Ca atoms in the solved protein structure
<sup>3</sup> Percentage of atoms that were predicted further than 3 Å away from any atoms in the solved structure
<sup>4</sup> Percentage of atoms from the solved structure that lay within 1 Å of a predicted atom
<sup>5</sup> Percentage of atoms from the solved structure for which the U-Net correctly predicted the secondary structure
<sup>6</sup> Percentage of atoms from the solved structure for which the U-Net correctly predicted the amino acid type (missing entries indicate that the solved structure did not contain amino acid type information)

In order to get a better idea of the implications of the results we take a closer look at the protein structures predicted by the DeepTracer and Phenix for the EMD-6272 density map (rotavirus VP6) and compare them to the solved structure (see **Fig 9**). We can see in the highlighted areas that the DeepTracer incorrectly connected amino acids in two cases. However, all predicted amino acids are still connected as a single chain. In comparison to that the prediction of Phenix is fragmented into multiple chains. Furthermore, amino acids appear to be missing from the prediction explaining the coverage difference of 8.1% (see Table 1). The predicted secondary structure elements in the Phenix prediction, particularly for sheets, appear also much less accurate than the ones made by the DeepTracer. Another noteworthy observation, although not a fair comparison as Phenix predicts more than just the backbone structure, is that the prediction runtime of Phenix for the EMD-6272 map was 1 hour and 45 minutes while the runtime of the DeepTracer for the same map was 40 seconds.

**Fig 9.**
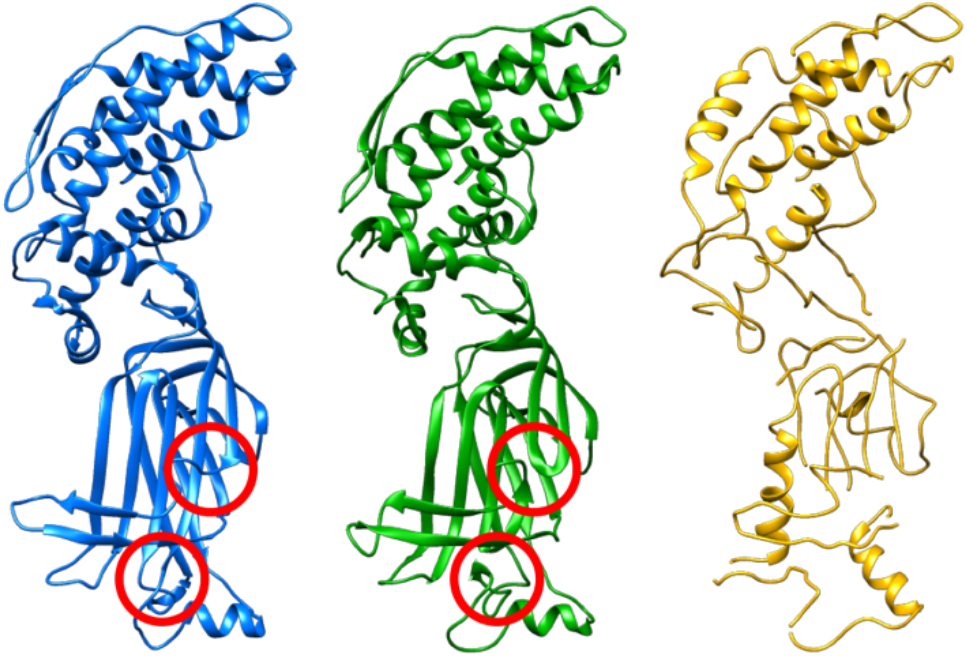
Solved PDB-3j9s protein structure (blue) and predictions by the DeepTracer (green) and Phenix (yellow) for EMD-6272 density map (rotavirus VP6). Highlighted areas show where the DeepTracer connected the amino acids incorrectly.

As mentioned in the introduction, a central goal of the DeepTracer was to improve the runtime of protein structure predictions. This makes it possible to predict ever larger density maps which allows for a more versatile application of the method. In order to compare the runtime of the DeepTracer with the C-CNN method we measure the execution time of each method for the prediction of multiple density maps. A scatter plot of the measured times is depicted in Fig 10.

**Fig 10.**
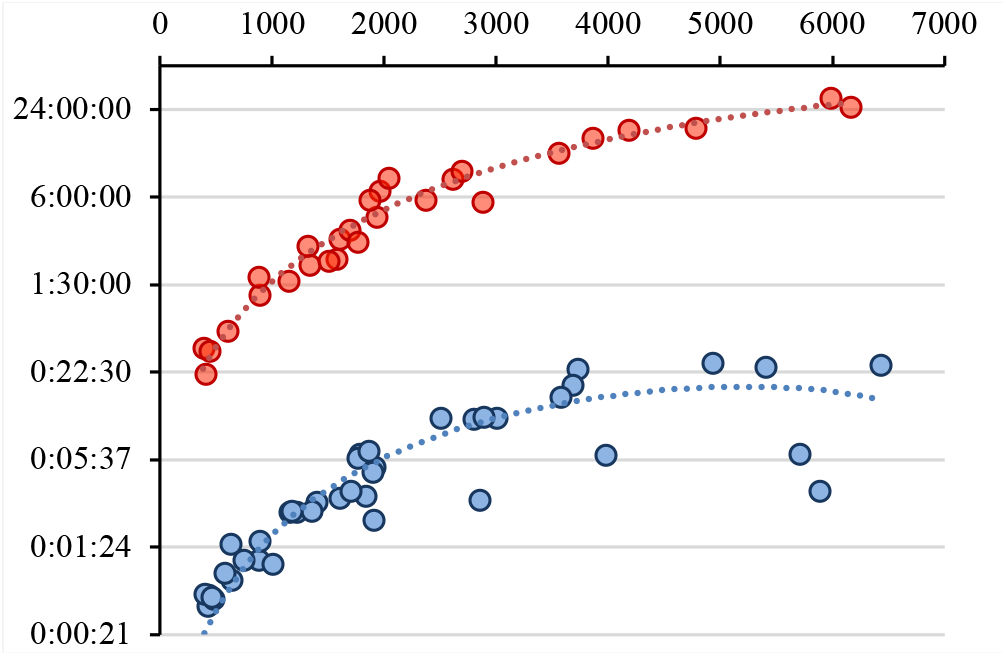
Scatter plot of prediction runtimes (y-axis) of the DeepTracer (blue) and the C-CNN method (red) for various density maps against the number of predicted atoms (x-axis). In order to see the runtimes more clearly, we use a logarithmic scale with a base of 4.

In order to evaluate the runtime of the DeepTracer for larger proteins we predicted the EMD-4903 density map (echovirus 1 intact particle). The number of predicted atoms for this map was 32,446. The execution of the end-to-end prediction pipeline of the DeepTracer took only two hours. As a point of comparison, the expected runtime of the C-CNN method for the same density map is more than 5 days assuming the trend in Fig 11 continues for larger proteins. The original density map next to the prediction of the DeepTracer can be seen in Fig 11.

**Fig 11.**
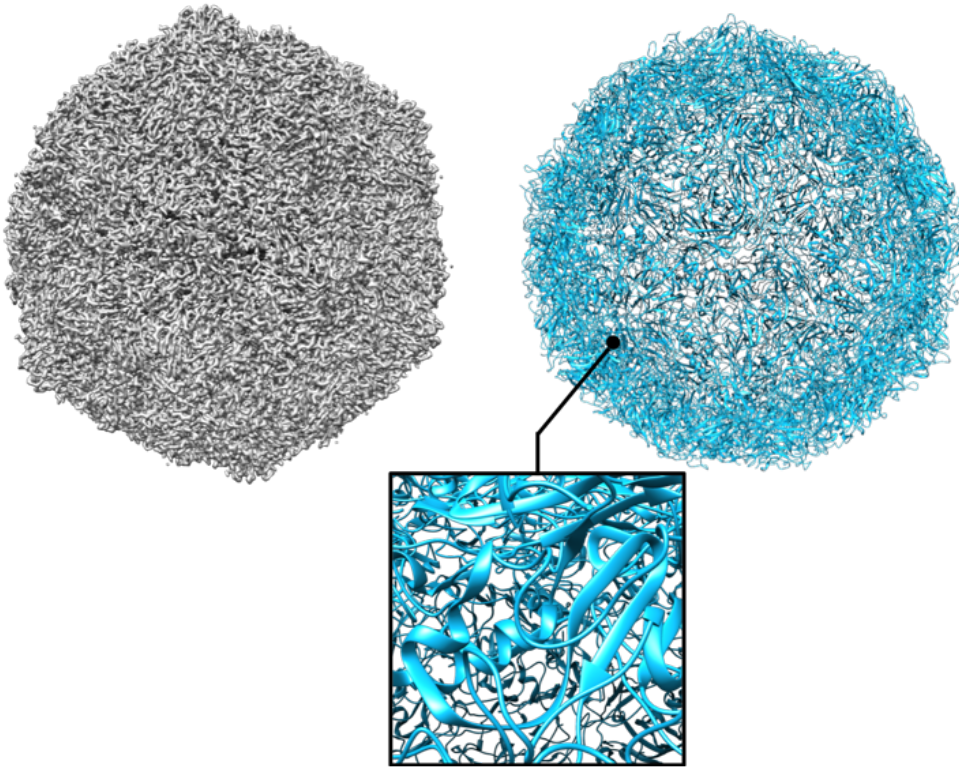
Density map (left) of EMD-4903 (echovirus 1 intact particle) and its predicted protein structure (right), containing more than 30,000 atoms.

In addition to the location and secondary structure of amino acids, the DeepTracer also predicts their types. As reported in **Table 1** we assigned the correct type out of the 20 possible amino acid types with an accuracy of 25.2%. In order to get a better idea of what that result means we look at a specific structure prediction. For the EMD-6272 density map, we predicted the correct amino acid type for 31.49% of amino acids. Note that this is predicted solely from 3D map without using any genetic information of the protein. In **Fig 12** we can see an extract of the amino acid sequence as predicted by the DeepTracer, compared to the sequence of the solved structure. The sequence was created by walking along the predicted chain of amino acids and noting down the type of each amino acid. The predicted amino acids from 3D could potentially be utilized for the reference of future full-atom structure prediction.

**Fig 12.**
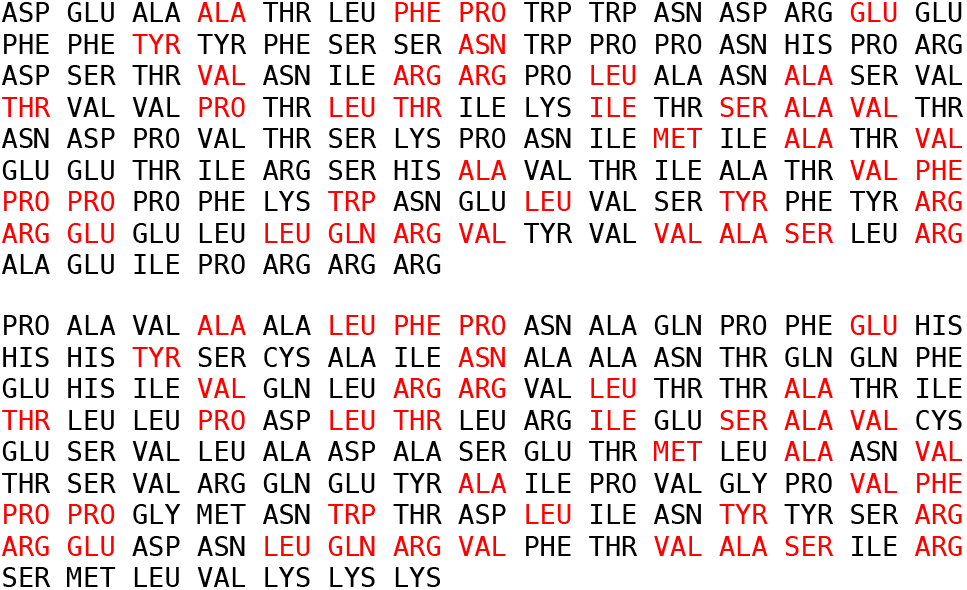
Extract of the amino acid sequence predicted by the DeepTracer for the EMD-6272 density map (top) compared to the sequence of the solved structure PDB-3j9s (bottom). Correctly predicted amino acids are highlighted in red.

## 3 Discussion

The final goal of the DeepTracer is to accurately predict the complete protein structure from high resolution cryo-EM density maps. In this section we evaluate how close we are to this goal, based on the results presented in the previous section, as well as discuss what further work is necessary.

When looking at the results reported in Table 1, we can note a couple of things. First, we see that the percentage of false positives predicted by the DeepTracer has increased by 1.9% compared to the C-CNN method. This can be explained through the manual thresholding which reduces noises in the experimental density maps and is utilized in the C-CNN method. However, considering that this step introduces manual effort to the prediction process, this is an acceptable concession as our new method is fully automated. Next, we see that the coverage percentage of the DeepTracer beats Phenix by over 20%. However, it still lacks behind the C-CNN method by 3%. On the contrary, Phenix performs best for RMSD, beating the DeepTracer by 0.08 and the C-CNN method by 0.09. If we compare the DeepTracer to Phenix based on these two metrics, we can see that the coverage improvements significantly outweigh the small deterioration in RMSD. However, compared to the C-CNN method there is no significant difference regarding the RMSD and coverage. As mentioned in the introduction, we wanted to focus not only on the prediction accuracy, but also on the runtime of the predictions. And here we can discern immense differences. For the prediction of smaller structures. the C-CNN method took at least 20 minutes while the DeepTracer ran small predictions in a matter of seconds. However, the bigger implications lay in the prediction of larger proteins. For around 6000 atoms the C-CNN method took over 24 hours to finish, whereas the DeepTracer only needed around 20 minutes, for some even less than 5 minutes. Particularly, the trendline is interesting as it gives us an idea of how the methods will behave for very large proteins. The C-CNN trendline seems almost linear above 2000 predicted proteins and the runtime seems to quadruple when 6000 predicted atoms are reached. If this trend continues then we could expect a 75-fold runtime increase in order to predict 100,000 atoms. The runtimes of DeepTracer, however, seem to remain flat once a certain number of atoms is reached which would mean that very large proteins could be predicted within a reasonable runtime. This can be explained by the chain identification step that is applied before the prediction of any atom. While larger proteins consist of more chains than smaller ones, the number of atoms in the chains does not necessarily grow. As each chain is processed separately this means that if the chain lengths remain constant and only the number of chains gets larger the runtime grows linearly with the number of chains.

In addition to the false-positive percentage, the coverage, and the RMSD we also calculated the secondary structure and amino acid type accuracy for the DeepTracer using the confidence maps predicted by U-Net. We can see that the secondary structure prediction performs well with an average accuracy of 81.72%. We also show that deep learning could discern different types of amino acids based on the density maps. Particularly, as this is only the preliminary stage of predicting amino acid type directly from 3D cryo-EM density map. It gives promise for future improvements through a more elaborate training of the U-Net with more data. Furthermore, we plan to include the amino acid sequence as an input parameter, so that we can map the predicted sequence with the input sequence and refine our prediction. For this sequence mapping work, an amino acid type prediction accuracy of around 30% from 3D could be already helpful. Particularly, as we can see from **Fig 12** that the U-Net tends to correctly predict the type of consecutive amino acids, likely due to high local resolutions. As it is very unlikely that this happens by chance, we can use this information to map the input sequence to our 3D backbone atom and amino acids prediction.

## 4 Conclusion

In this paper, we presented the DeepTracer, a fully automated tool which can accurately predict backbone protein structures based on 3D cryo-EM density maps. In terms of accuracy, we were able to outperform the established structure prediction method Phenix by over 20% in structure coverage with only a slight RMSD increase of 0.08. Compared to the deep learning-based C-CNN method we achieved a slightly better RMSD by a margin of 0.01 with a slightly decreased coverage of 3%. In contrast with the C-CNN method, this fully automated method requires no manually set parameters. And the vastly reduced runtime allows us to predict protein structures within minutes instead of days. This is significant, as it allows the researchers to apply the DeepTracer on very large density maps and expect results maximally within a couple of hours. Furthermore, we achieved preliminary results in the amino acid type prediction from 3D with an accuracy of 25.2%. This gives promise to the incorporation of amino acid sequence into the prediction pipeline. Since the predicted amino acids from 3D could potentially be utilized for the reference of future full-atom structure prediction. In the future, we expect to use the amino acid type information to extend the functionality of the DeepTracer such that it can predict the protein full atom structure, including the side chains of the amino acids.

## Footnotes

1 https://docs.scipy.org/doc/scipy/reference/generated/scipy.optimize.minimize.html

